# BRUCE liver-deficiency potentiates MASLD/MASH in PTEN liver-deficient background by impairment of mitochondrial metabolism in hepatocytes and activation of STAT3 signaling in hepatic stellate cells

**DOI:** 10.1101/2024.09.13.611500

**Authors:** Lixiao Che, Camille K. Stevenson, David R. Plas, Jiang Wang, Chunying Du

## Abstract

Metabolic dysfunction-associated steatotic liver disease (MASLD) is currently the most common liver disease, affecting up to 25% of people worldwide, featuring excessive fat accumulation in hepatocytes. Its advanced form, metabolic dysfunction-associated steatohepatitis (MASH), is a serious disease with hepatic inflammation and fibrosis, increasing the need for liver transplants. However, the pathogenic mechanism of MASLD and MASH is not fully understood. We reported that BRUCE (*BIRC6)* is a liver cancer suppressor and is downregulated in MASLD/MASH patient liver specimens, though the functional role of BRUCE in MASLD/MASH remains to be elucidated. To this end, we generated liver-specific double KO (DKO) mice of BRUCE and PTEN, a major tumor suppressor and MASLD/MASH suppressor. By comparing liver histopathology among 2-3-month-old mice, there were no signs of MASLD or MASH in BRUCE liver-KO mice and only onset of steatosis in PTEN liver-KO mice. Interestingly, DKO mice had developed robust hepatic steatosis with inflammation and fibrosis. Further analysis of mitochondrial function with primary hepatocytes found moderate reduction of mitochondrial respiration, ATP production and fatty acid oxidation in BRUCE KO and the greatest reduction in DKO hepatocytes. Moreover, aberrant activation of pro-fibrotic STAT3 signaling was found in hepatic stellate cells (HSCs) in DKO mice which was prevented by administered STAT3-specific inhibitor (TTI-101). Collectively, the data demonstrates by maintaining mitochondrial metabolism BRUCE works in concert with PTEN to suppress the pro-fibrogenic STAT3 activation in HSCs and consequentially prevent MASLD/MASH. The findings highlight BRUCE being a new co-suppressor of MASLD/MASH.

## Introduction

Fatty liver disease, now called metabolic dysfunction-associated steatotic liver disease (MASLD), is a global epidemic that is estimated to affect up to one-quarter of the world’s population and occurs when excessive fat is accumulated in hepatocytes. MASLD can progress to the more severe form of metabolic dysfunction-associated steatohepatitis (MASH) with features of hepatic inflammation, hepatocyte ballooning and fibrosis. Importantly, hepatic stellate cells (HSCs) play a crucial role in the development of liver fibrosis by secreting fibrogenic factors that promote liver fibrosis. MASH can lead to cirrhosis of the liver with tissue scarring; hence it is becoming a common indication for liver transplants. It is estimated as many as 15 million people in the United States are affected by MASH; however, the mechanisms underlying the pathogenesis of MASLD, and MASLD-to-MASH progression are still not fully understood. Therefore, discovery of new pathogenic mechanisms underlying both conditions is of paramount importance^1^.

MASLD is the consequence of lipid metabolic dysregulation. Mitochondria play critical roles in both lipid metabolism and bioenergetics via fatty acid β-oxidation (FAO) and respiration/oxidative phosphorylation (OXPHOS), respectively. Mitochondrial FAO is a major fatty acid catabolism pathway that facilitates lipid clearance for lipid homeostasis. Mitochondrial respiration uses the electron transport chain to produce ATP from macromolecules through OXPHOS. Clinically, dysfunction of both mitochondrial FAO and bioenergetics is closely associated with MASLD and MASH. Studies with MASLD/MASH patient liver specimens found up to a 50% reduction of the capacity of mitochondrial FAO compared to healthy subjects with normal histology^2^. In addition, liver tissues from MASLD and MASH specimens have ultrastructural mitochondrial damage, reduced FAO and respiratory chain activity, ATP depletion, and reactive oxygen species (ROS) overproduction^3,4^. Despite the clinical link of mitochondrial dysfunction to MASLD/MASH in humans, the protein factors that lead to MASLD/MASH development through dysregulation of mitochondrial metabolism are not fully understood. Further, only a portion of patients with MASLD will develop MASH, and the determinants in this partial progression are not fully elucidated.

Outside of mitochondria, dysregulation of various metabolic pathways could result in MASLD/MASH and progression to hepatocellular carcinoma (HCC). Among these, the dysregulation of the phosphoinositide 3-kinase (PI3K)/ phosphatase and tensin homolog (PTEN)/AKT (i.e., PI3K-PTEN-AKT) pathway contributes to MASLD and MASH leading to HCC and intrahepatic cholangiocarcinoma (ICC)^5^. PI3K phosphorylates the substrate phosphatidylinositol 4,5-biphosphate (PIP2) to form phosphatidylinositol 3,4,5-triphosphate (PIP3) on intracellular membranes, subsequently recruiting signaling proteins of AKT and others leading to lipogenesis and promotion of HCC development^6^. In contrast to PI3K, PTEN, a potent tumor suppressor, has a phosphoinositide phosphatase activity. By dephosphorylating PIP3 to PIP2, PTEN switches off the PI3K-AKT signaling axis and represses PIP3-mediated activation of AKT and hence AKT-mediated cellular functions of lipogenesis^7^. In MASLD patients, reduced expression of PTEN and concurrent hyperactivation of AKT are found in liver biopsy tissue, suggesting hepatic steatosis can be mediated by reduced PTEN expression in hepatocytes^8^. Studies have shown liver-specific PTEN-KO in mice de-represses the PI3K-AKT pathway and thereby promotes lipogenesis with a MASLD onset at 2–3 months of age^5,9^ and mild fibrosis in some mutant livers around one year of age^5^. When mice age further, PTEN liver-deficiency promotes cell growth and proliferation leading to HCC and ICC starting at 12-months-old^9^. Although the function of PTEN in suppressing liver tumorigenesis is well appreciated, how PTEN represses MASLD/MASH and which other hepatic proteins work in concert with PTEN in the suppression of the pre-cancerous MASLD/MASH is less clear.

BIR repeat-containing ubiquitin-conjugating enzyme (BRUCE) is a hybrid ubiquitin conjugase and ligase and a scaffolding protein. Originally, BRUCE was found as a member of the inhibitor of apoptosis protein (IAP) family^10^. Knockdown of BRUCE sensitizes tumor cells to apoptosis^11,12^ as BRUCE ubiquitinates various pro-apoptotic proteins for their degradation via the ubiquitin-proteasomal system^13^. We have reported that BRUCE whole-body mutant mice have activation of p53-dependent apoptosis pathway and embryonic lethality^14^. Since then, the versatility of BRUCE function has begun to unfold outside of its role in anti-apoptosis. We reported that BRUCE is a crucial stabilizer for genomic stability and a suppressor of liver fibrosis and tumorigenesis induced by hepatocarcinogen DEN^15,16^. We also reported BRUCE is required for the maintenance of germ cell genome integrity via repair of physiological DNA breaks during spermatogenesis in male mice^17^. The diversity of BRUCE function is further shown by our published study reporting the role of BRUCE in autophagy-regulated cellular energy metabolism in human tumor cell lines. Knockdown of BRUCE reduces cellular energy levels (ATP) and induces autophagy by driving activation of the AMPK-ULK1 autophagic initiating axis. So, BRUCE-KD cells use autophagy as a means to increase energy supply in human tumor cells^18^. As BRUCE liver-KO young mice do not develop spontaneous diseases or tumors but significantly exacerbates hepatocarcinogen-initiated liver fibrosis and HCC, we propose BRUCE acts as a modifier of liver pathogenesis that works in concert with other liver disease drivers.

Clinically, we have found that BRUCE is downregulated in 50% of MASLD, MASH and 84% of HCC specimens^15^. However, evidence of BRUCE function in the etiology of MASLD/MASH is lacking. We sought to examine whether BRUCE works in concert with PTEN to co-regulate MASLD/MASH development. In this study, we provided evidence for BRUCE being a crucial co-regulator working in concert with PTEN to suppress hepatosteatosis and liver fibrosis.

## Results

### 1. BRUCE liver-KO exacerbates MASLD in PTEN liver-deficient background

Young adult BRUCE liver-KO (BKO) mice (2–3-months-old) did not show noticeable hepatic histological and pathological changes, so single BRUCE-deficiency is insufficient to induce spontaneous liver disease in young mice^15^. BRUCE liver-KO mice displayed prominent exacerbation of hepatic fibrosis and HCC development under the tumorigenic condition of DEN exposure^15^. In that model, lipid accumulation in hepatocytes was observed, suggesting that BRUCE liver-KO may influence steatosis and steatohepatitis. Therefore, we postulated the impact of BRUCE liver-KO on MASLD/MASH might be revealed under a stressed, MASLD- and MASH-inducing condition. To test this possibility, we selected the PTEN liver-KO (PKO) mouse model as the MASLD/MASH background, because it is an established model, starting MASLD in young adults (3-months-old) and further progressing to MASH starting after 6 months of age^5,9^. We bred BRUCE liver-deficient (*Bruce^fl/fl^AlbCre*^+^ BKO) mice^15^ into the background of PTEN liver-deficiency (*Pten^fl/fl^AlbCre*^+^ PKO). The resultant four cohorts of mice, ‘WT’ (*AlbCre^-^),* PKO, BKO and Double KO (DKO; *Bruce^fl/fl^Pten^fl/fl^AlbCre*^+^), were genotypically validated by PCR of genomic DNA (**Fig. S1**). Further, ablation of PTEN and BRUCE protein expression was validated by Western blotting (WB) with whole liver tissue homogenates (**Fig. 1A**). The WB results show that deficiency of either one did not influence the expression of the other in the liver, indicating that a confounding effect from loss of one protein causing loss of the other is not likely to be an issue.

**Figure 1.**
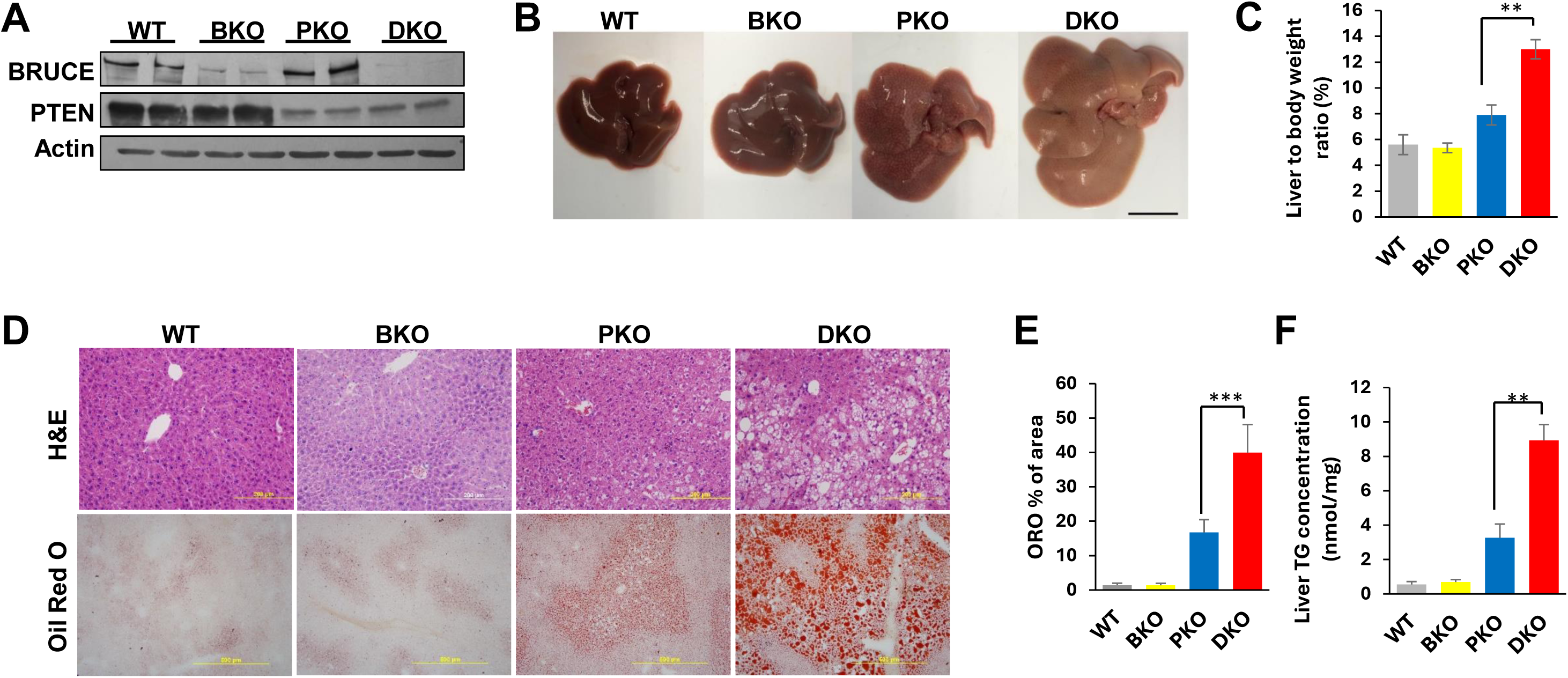
BRUCE liver-KO accelerates liver MASLD in PTEN KO background. **(A)** liver homogenates prepared from the four genotypes of mice were analyzed by Western Blotting (WB) for expression of BRUCE, PTEN and Actin (control) using an antibody specific to each protein. **(B)** macroscopic liver images of the four genotypes of mice (scale bar, 1cm). **(C)** quantification of the liver to body weight ratio of the indicated mice. **(D)** H&E analysis of liver tissue sections (upper) and Oil Red O staining of hepatic lipid (lower) of indicated mice. **(E)** quantification of Oil Red O (ORO) results. **(F)** quantification of total liver triglyceride (TG) in four genotypes of mice. (n.s.: not significant, **p<0.01, ***p<0.001 by student’s *t* test).

To examine the effect of BRUCE liver-deficiency on the development of MASLD and MASH in PTEN-KO background, we conducted a time course study to compare the liver phenotypic changes among the four genotypes. At 2-3 months of age, BKO mice showed no noticeable liver alterations, and their livers were comparable to WT livers in size and color (**Fig. 1B**) and in liver to body weight ratio (**Fig. 1C**). However, livers from DKO mice were bigger and paler with a higher liver-to-body weight ratio than the livers from PKO (**Fig. 1B and 1C**). These results indicate that although BRUCE liver-KO mice have not shown noticeable liver phenotypes, BRUCE deficiency has imposed an exacerbating effect on hepatic lipid accumulation in PTEN liver-KO background (i.e., DKO). Indeed, H&E staining of liver tissue sections showed a mild lipid droplet increase in PKO, and an excessive lipid accumulation in DKO mice with mixed macro- and micro-steatosis and associated hepatocyte ballooning (**Fig. 1D**). The lipid accumulation was further verified by Oil Red O (ORO) staining of hepatic neutral lipids (**Fig. 1D**) and by quantification of the ORO staining results (**Fig. 1E**). Further, the lipid accumulation was validated by increased total liver triglyceride (TG) contents (**Fig. 1F**). Collectively, these data show a contribution of BRUCE liver-deficiency in the PTEN liver-deficient background by imposing a significant exacerbating effect on MASLD as early as two months of age, when BRUCE liver-deficiency does not display noticeable MASLD. Since PKO and DKO mice at 2-3-months-old just begin to show MASLD, investigation into the pathological mechanism at this beginning stage could reveal the initial driving event(s) underlying the exacerbating function by BRUCE liver-deficiency on PTEN KO background. Therefore, mice at 2-3 months of age are the subjects of this study.

### 2. BRUCE liver-deficiency attenuates mitochondrial fatty acid β-oxidation which contributes to the exacerbated steatosis in PTEN-deficient background

Loss of PTEN de-represses the AKT-PI3K pathway and promotes de novo lipogenesis, which is a major cause of hepatosteatosis induced by PTEN liver-deficiency^19–21^. To investigate how BRUCE liver-deficiency exacerbates MASLD in PTEN KO background, we first examined whether BRUCE liver-deficiency further increases pro-steatotic AKT signaling. By WB analysis of hepatic pro-lipogenic AKT signaling components with whole liver tissue homogenates, an expected AKT activation was observed in PTEN KO liver, as indicated by phosphorylation of AKT at Ser-473, which was further validated by phosphorylation of a protein substrate of AKT, GSK3β at Ser-9 (**Fig. 2A**, PKO vs WT). However, no increase in the levels of total AKT or pAKT-Ser473, as well as total GSK3β or pGSK3β-Ser9 were observed in BKO mice and no further increase of them in DKO compared to PKO (**Fig. 2A**). Furthermore, at lipogenic gene expression levels, BKO did not display elevated expression of key lipogenic genes *Fasn* and *Acly* than WT control, nor DKO showed elevated expression of these genes than PKO (**Fig. S2A).** Together the data suggests BRUCE liver-deficiency does not elevate AKT-mediated lipogenesis and as such, the BKO-exacerbated MASLD in PKO background is through other pathways than AKT-mediated lipogenesis.

**Figure 2.**
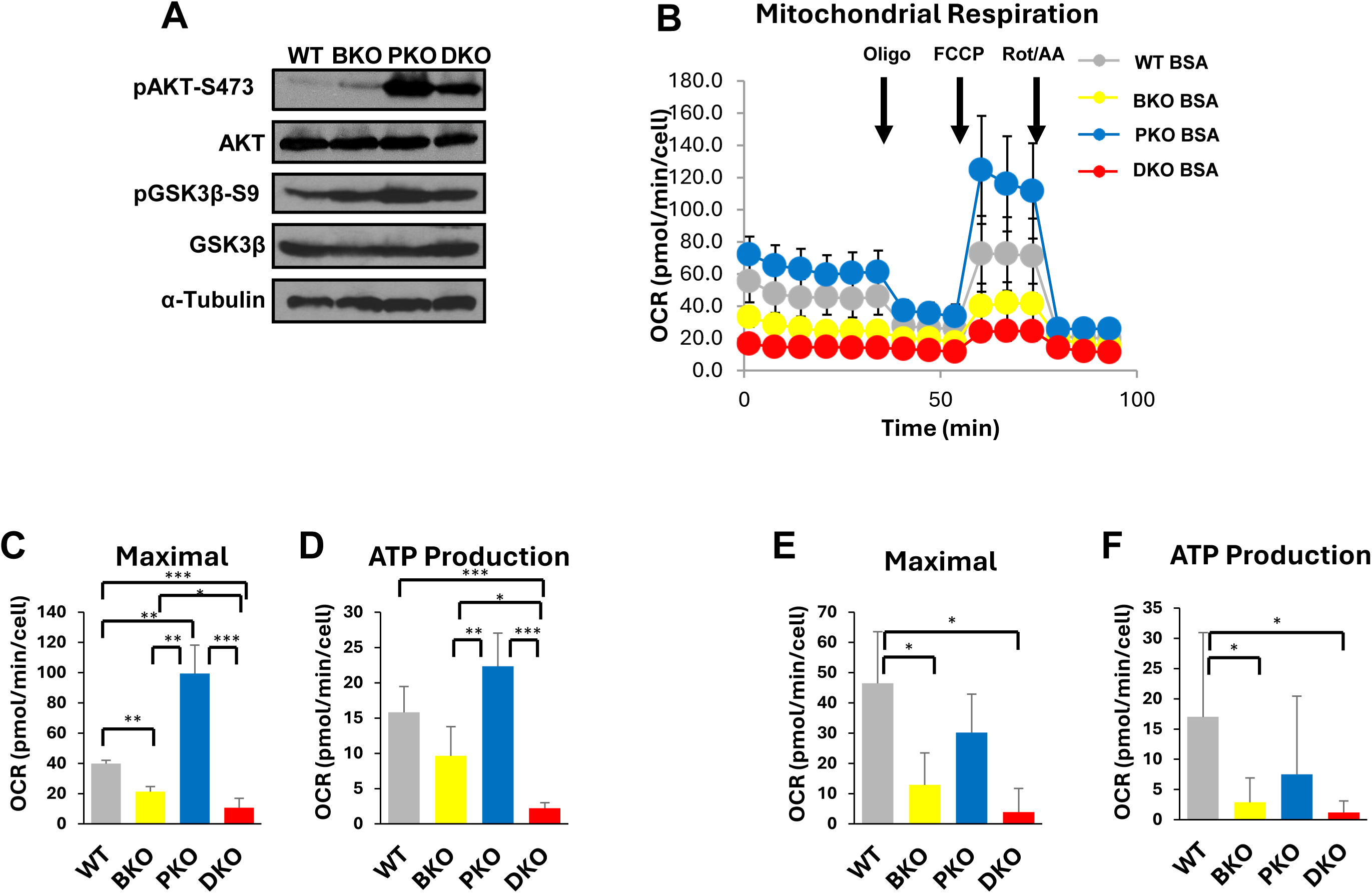
BRUCE liver-KO hepatocytes exhibit defects in mitochondrial metabolism. **(A)** WB analysis of (p)AKT, (p)GSK3β, and Tubulin (control) in total liver homogenates from mice of each genotype. **(B)** kinetics of mitochondrial respiration shown by oxygen consumption rate (OCR) measured in primary hepatocytes after sequential compound injections of oligomycin (Oligo), carbonyl cyanide-p-trifluoromethoxyphenylhydrazone (FCCP), and rotenone + Antimycin A (Rot/AA). **(C)** quantification highlighting differences in the mitochondrial functions of maximal respiration and **(D)** ATP production between genotypes. To measure respiration due to the use of exogenous fatty acids, primary hepatocytes were treated with palmitic acid (PA) for 24 hours and etomoxir (ETO) injections were added prior to analysis; the PA treated values minus PA + ETO treated values resulted in the graphs for maximal respiration **(E)** and ATP production supported by FAO **(F)**. (*p<0.05, **p<0.01, ***p<0.001 by student’s *t* test).

In addition to AKT-PI3K pro-steatotic pathway, mitochondria play a crucial role in the regulation of MASLD development. Reduced mitochondrial FAO, respiration and bioenergetics (ATP production) are tightly associated with MASLD patients^22^. Therefore, we investigated whether BRUCE liver-deficiency impacts these mitochondrial functions in fresh hepatocytes isolated from livers of the four genotypes of mice using a Seahorse analyzer. Hepatocytes were seeded in Seahorse XF cell culture plates, treated with specific mitochondrial respiration complex inhibitors in a temporal manner as indicated (**Fig. 2B**), and the mitochondrial basal and maximal respiration, FAO, and ATP production were measured in real time in these cells. The results showed that both the basal and maximal respiration were reduced in BKO hepatocytes compared to WT while reduction of ATP generation was not significant (**Figs. S2C, 2C-D**), suggesting that BRUCE is required for a proficient mitochondrial respiration. Remarkably, both processes were greatly reduced in DKO cells (**Fig. 2C-D**), indicating that BRUCE is required to support mitochondrial respiration and ATP production in PTEN-KO background.

Mitochondrial FAO is the major fatty acid catabolic pathway essential for lipid homeostasis by reducing excessive lipids. As such, impaired mitochondrial FAO is one of the major clinical features in patients with MASLD and MASH^2^. Using these hepatocytes, we measured mitochondrial oxidation of the long-chain fatty acid, palmitic acid (PA). Hepatocytes were treated with BSA (control) or 100 µM PA for 24 hours, followed by acute injection of etomoxir (ETO) prior to measurement of mitochondrial function. Etomoxir is an irreversible inhibitor of carnitine palmitoyl transferase 1a (CPT1a) which transports exogenous fatty acids into mitochondria for oxidation. Hence, CPT1a inhibition prevents entry of new fatty acids into mitochondria. Compared to WT hepatocytes, BKO, but not PKO cells, displayed decreased FAO (the value of PA treatment minus that of PA+ETO), indicating BRUCE is required for mitochondrial oxidation of long-chain fatty acids. Remarkably, among the four cell types, the lowest oxidation was observed in DKO hepatocytes (**Figs. S2E, 2E-F)**, indicating BRUCE and PTEN work in concert to maintain efficient fatty acid oxidation for lipid clearance. Collectively, these results support a new function of hepatic BRUCE in the maintenance of mitochondrial regulation of lipid homeostasis, at least by promotion of mitochondrial FAO and bioenergetics in the face of higher lipid levels in PTEN-KO background.

Hepatic lipid accumulation is driven by the dysregulation of lipid metabolism. The data indicate that both PTEN and BRUCE each as a single agent have maintenance roles in lipid homeostasis and their dual deficiencies contribute to accelerated MASLD progression via aberrant AKT-lipogenesis activity and mitochondrial metabolic defects, respectively. At 2-3 months, single KO of PTEN promotes MASLD onset due to elevated lipogenesis, yet FAO is maintained due to the presence of BRUCE. While BRUCE single KO impairs FAO and fat clearance, PTEN represses AKT-lipogenesis, preventing MASLD development. There is a dynamic balance between lipid production and catabolism to maintain homeostasis and avert hepatic steatosis. In BRUCE and PTEN DKO mice, there is no compensation by either protein, driving MASLD due to the combination of attenuated mitochondrial metabolic capacity and overproduction of lipids from overactivated PI3K-AKT-lipogenesis. This supports why DKO mice exhibit accelerated MASLD compared to PKO.

### 3. BRUCE liver-deficiency exacerbates both oxidative stress and liver injury in PTEN liver-KO background

Given the tight association of dysregulated mitochondrial lipid metabolism and bioenergetics with both oxidative stress and liver injury in MASLD and MASH, we assessed the levels of superoxide anions as an indicator of oxidative stress by a superoxide indicator dihydroethidium (DHE) in the four genotypes of mouse livers. DHE exhibits blue fluorescence in the cytosol until oxidized, where it intercalates within the cell’s DNA and stains its nucleus a bright fluorescent red. We found that both WT and BKO liver tissue sections did not show noticeable nuclear DHE fluorescence; however, a higher level of fluorescence was observed in PKO, and further augmented in DKO liver tissue sections (**Fig. 3A**). To evaluate whether the oxidative stress is associated with liver injury, we examined DNA damage and apoptosis because BRUCE has roles in promotion of DNA strand break repair and inhibition of apoptosis^14,23,24^. In agreement with no elevation in oxidative stress in BKO liver, neither DNA strand breaks assessed by IHC of γH2AX, nor apoptosis assessed by both IHC of cleaved caspase-3 and TUNEL were detected in BKO livers (**Fig. 3B-D**), suggesting BRUCE liver-deficiency alone does not induce spontaneous DNA damage or apoptosis in the absence of hepatic stress. However, under the steatotic stress induced by PTEN-KO background, BRUCE liver-deficiency further elevated liver injury as displayed in the DKO liver (**Fig. 3B-D**). Apoptosis-induced loss of hepatocytes can be replenished by new hepatocytes via compensatory proliferation which is another indicator of liver injury^25^. An increase in hepatocyte compensatory proliferation indicated by Ki67 expression was found in the DKO liver (**Fig. 3E**). Collectively, BRUCE KO exacerbates oxidative stress and liver injury in PTEN-KO steatotic background.

**Figure 3.**
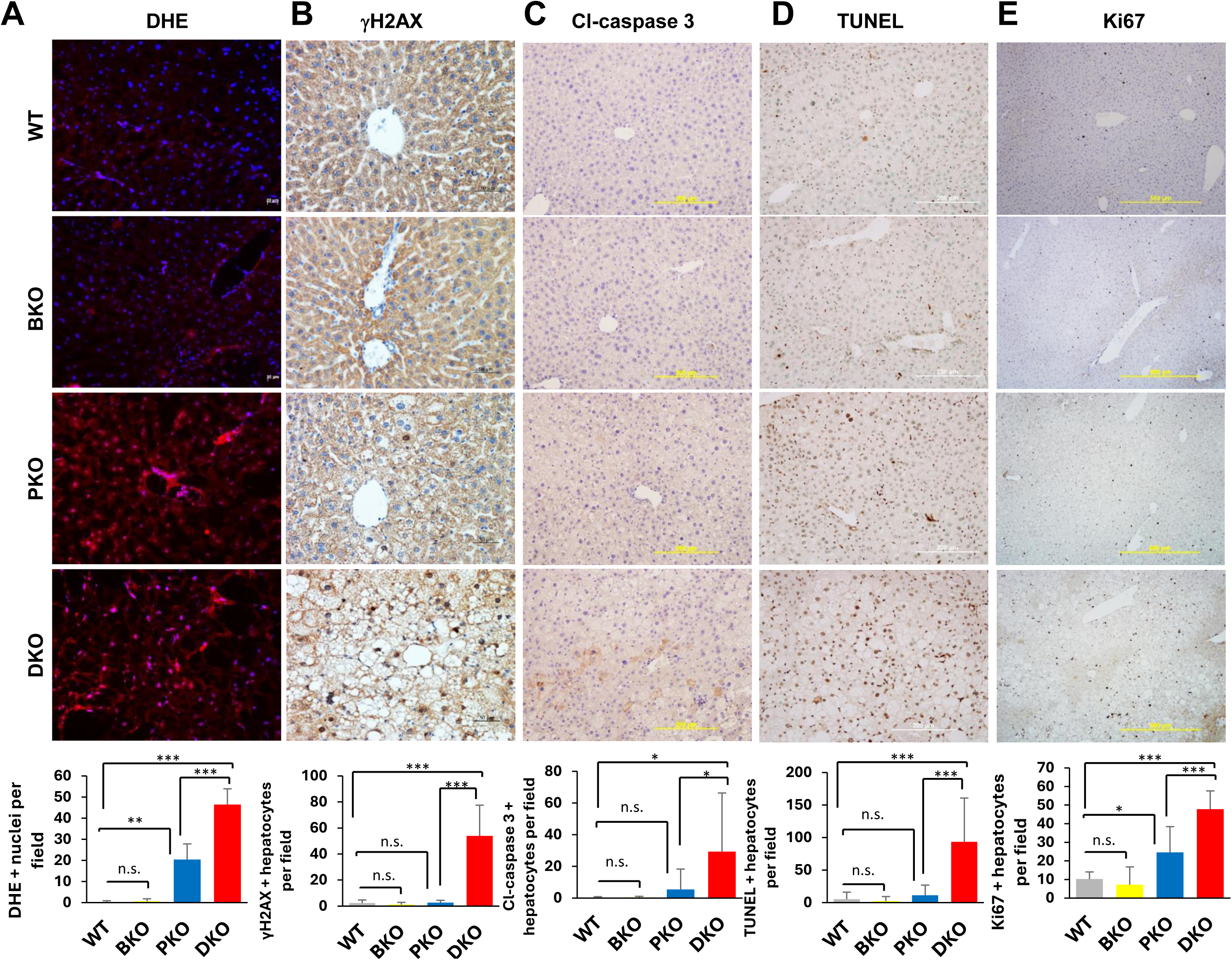
BRUCE liver-KO exacerbates oxidative stress and liver injury in PTEN KO background. Staining of liver sections from mice of each indicated genotype with dihydroethidium (DHE) **(A),** γH2AX **(B)**, cleaved caspase 3 (Cl-caspase 3) **(C)**, TUNEL **(D)**, and Ki67 **(E)**, followed by quantifications underneath. (n.s.: not significant, *p<0.05, **p<0.01, ***p<0.001 by student’s *t* test).

### 4. BRUCE liver-deficiency accelerates MASLD-to-MASH transition in PTEN-KO background

It is known that PKO mice begin to progress from MASLD to MASH around 10 months of age^5,9^. Given that oxidative stress and liver injury promote both proinflammatory and pro-fibrotic responses in liver^26^, we assessed whether BKO accelerates the transition from MASLD to MASH in 2-month-old DKO mice. Hepatic inflammatory response was assessed by the number of mature macrophages expressing F4/80 and the number of neutrophils by an NIMP-R14 antibody. We found no noticeable increase of both in BKO compared to WT liver, an increase in macrophages but not neutrophils in PKO liver, and significant increase of both in DKO mice (**Fig. 4A-B**). Remarkably, among the four genotypes, livers from DKO mice exhibited the most severe hepatic fibrosis by Sirius Red staining of collagen accumulation, and the most robust activation of hepatic stellate cells (HSCs) by α-SMA expression (**Fig. 4C-D**). The elevated activation of HSCs was further supported at gene expression level by the whole liver transcriptomics analysis showing a significant upregulation of three critical gene transcripts involved in HSC activation, *Cola1*, *S100A6* and *Timp1* (**Fig. 4E**). Collectively, although BRUCE liver-KO is insufficient to induce inflammation and fibrosis at 2-3-months-old, it accelerates the transition from MASLD-to-MASH in PKO liver.

**Figure 4.**
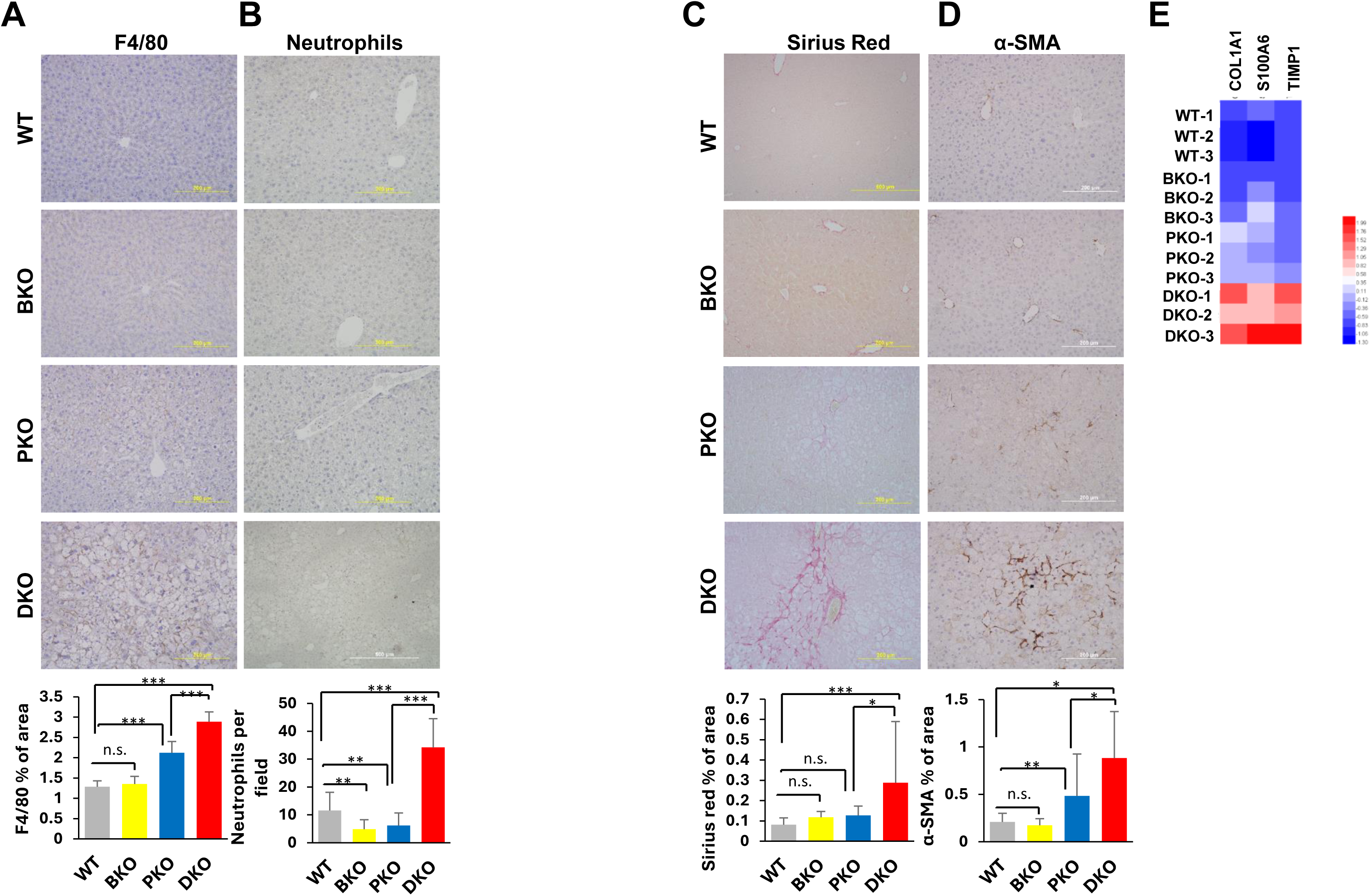
BRUCE liver-KO accelerates MASH progression in PTEN-KO background. **(A)** representative liver sections with F4/80 immunohistochemistry staining, quantified below. **(B)** neutrophil staining in each genotype with quantification below. Sirius Red **(C)** and α-SMA staining **(D)** across the indicated genotypes with the respective quantifications below. **(E)** heatmap of relative mRNA expression of hepatic stellate cell activation genes generated by RNA sequencing analysis. (n.s.: not significant, *p<0.05, **p<0.01, ***p<0.001 by student’s *t* test).

### 5. Molecular and cellular events that mediate MASLD-to-MASH transition in BRUCE and PTEN DKO mice

To understand the molecular and cellular mechanisms underlying the above inflammatory and fibrotic responses, we assessed four crucial events that promote hepatic inflammation and fibrosis. Liver injury by inflammation can cause expansion of bile ductal cells from activation of biliary/progenitor cells. By conducting IHC with an antibody against cytokeratin 19 (CK19), a marker of biliary/progenitor cells, we found the highest CK19-postive cell numbers in DKO livers suggesting the greatest expansion of bile ductal cells among all four samples (**Fig. 5A**). In addition, the transcription factor sex determining region Y box 9 (SOX9) becomes expressed during HSC activation by profibrotic signaling factors when it promotes production of extracellular matrix (ECM) components such as type 1 collagen (COL1) among others^27^. We found the highest expression level of SOX9 in DKO livers among the four samples (**Fig. 5B**). Further, DKO livers showed the greatest expression of Cluster of Differentiation 44 (CD44) (**Fig. 5C**), a cell surface marker protein mainly expressed in immune cells and overexpressed in activated HSCs and believed to play a key role in MASLD-to-MASH transition by regulating hepatic macrophage polarization (pro-inflammatory phenotype) and infiltration (macrophage motility and the MCP1/CCL2/CCR2 system)^28^ as well as active HSC-mediated fibrosis^29^. Moreover, DKO livers showed the highest level of β-catenin expression (**Fig. 5D**), a facilitator of both liver fibrosis and MASLD-to-MASH progression, and an accelerator of hepatocellular carcinoma^16,30–33^. Collectively, these pro-inflammatory and pro-fibrogenic events provide a critical molecular basis for the accelerated MASLD-to-MASH transition in DKO mice.

**Figure 5.**
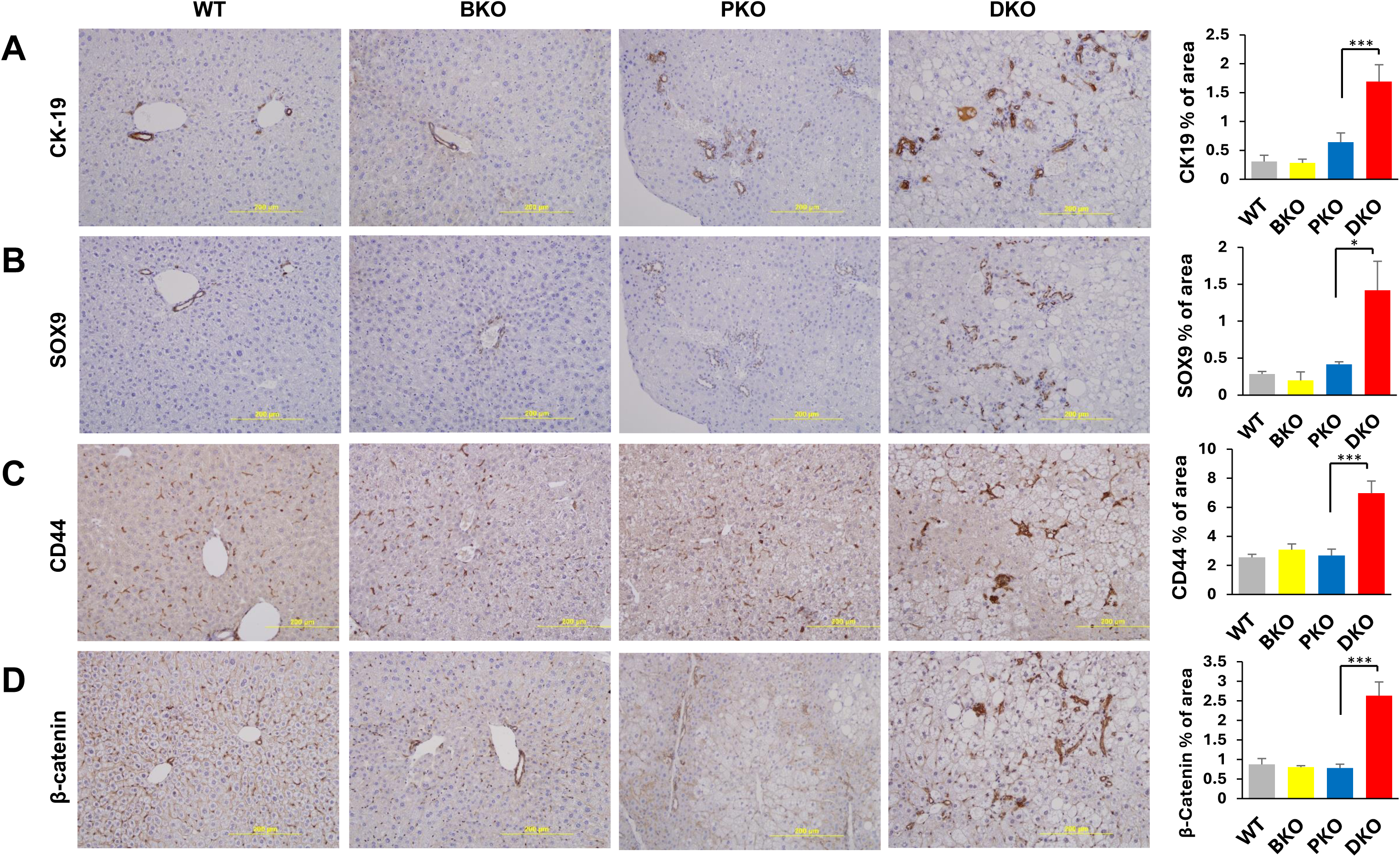
Molecular events that underscore inflammatory and fibrotic responses in BRUCE and PTEN double liver-KO mice. Immunohistochemistry staining of CK-19 **(A)**, SOX9 **(B)**, CD44 **(C)** and β-catenin **(D)** in liver sections from each genotype, all followed by quantification to the righthand side. (*p<0.05, ***p<0.001 by student’s *t* test).

### 6. Dual liver-deficiency of BRUCE and PTEN in hepatocytes activates pro-fibrotic STAT3 signaling in HSCs

To gain insights into how dual liver-deficiency of BRUCE and PTEN accelerates inflammation and fibrosis during the transition from MASLD to MASH, we analyzed our whole liver tissue transcriptomics data. By conducting gene expression analysis using the HemI software, we found a panel of significantly upregulated transcripts of proinflammation genes, cytokines and related receptors exclusively in DKO liver tissue, and among them was the signal transducer and activator of transcription 3 (*Stat3*) (**Fig. 6A**), which regulates a large repertoire of genes and is a key regulator of liver inflammation and fibrosis^34^. As a transcription factor, STAT3 is activated in the cytoplasm by phosphorylation in response to cytokines and growth factors under pathological conditions, then translocates to cell nucleus to activate gene expression. As phosphorylation at amino acid tyrosine 705 (Y705) is a key event in STAT3 activation^35,36^, we examined STAT3 activation by performing WB analysis with an antibody specific for pSTAT3-Y705 in whole liver homogenates. We found pSTAT3-Y705 was increased only in DKO liver tissue, and the total STAT3 protein levels remained unchanged across the four genotypes (**Fig. 6B**), demonstrating an exclusive STAT3 activation in DKO livers. In addition to parenchymal hepatocytes, liver tissue comprises a host of specialized non-parenchymal cells (NPCs), which critically participate in hepatic inflammation and fibrosis. To distinguish the cell type in which STAT3 is activated, we examined the expression of active, nuclear localized pSTAT3-Y705 and that of a cell-type marker by IHC performed on consecutive liver tissue slides to identify the cell type that expresses pSTAT3-Y705. We found that cells positive for nuclear expression of STAT3-Y705 were also positive for the expression of HSC marker α-SMA, hence these cells are activated HSCs (**Fig. 6C**). The data indicates that among multiple pathways activated by oxidative stress, STAT3 is activated specifically in the HSC cell population through its tyrosine phosphorylation and its nuclear translocation in BRUCE and PTEN DKO mice. HSCs play a key role in the development of MASH because activated HSCs are a major source of ECM production in liver fibrosis and responsible for liver inflammation induced by cytokines. Although activated STAT3 signaling in liver inflammation and fibrosis is known, whether STAT3 activation in HSCs impacts MASLD-to-MASH progression is not fully understood

**Figure 6.**
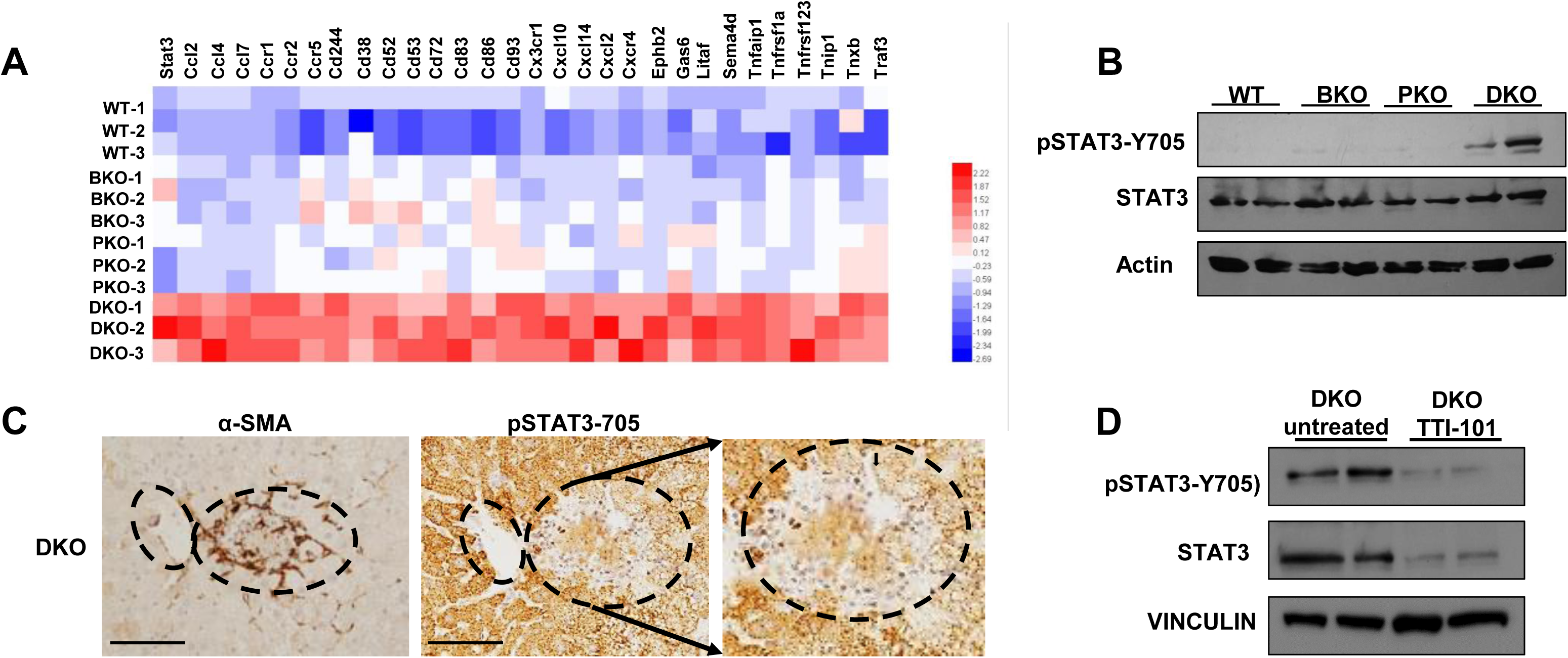
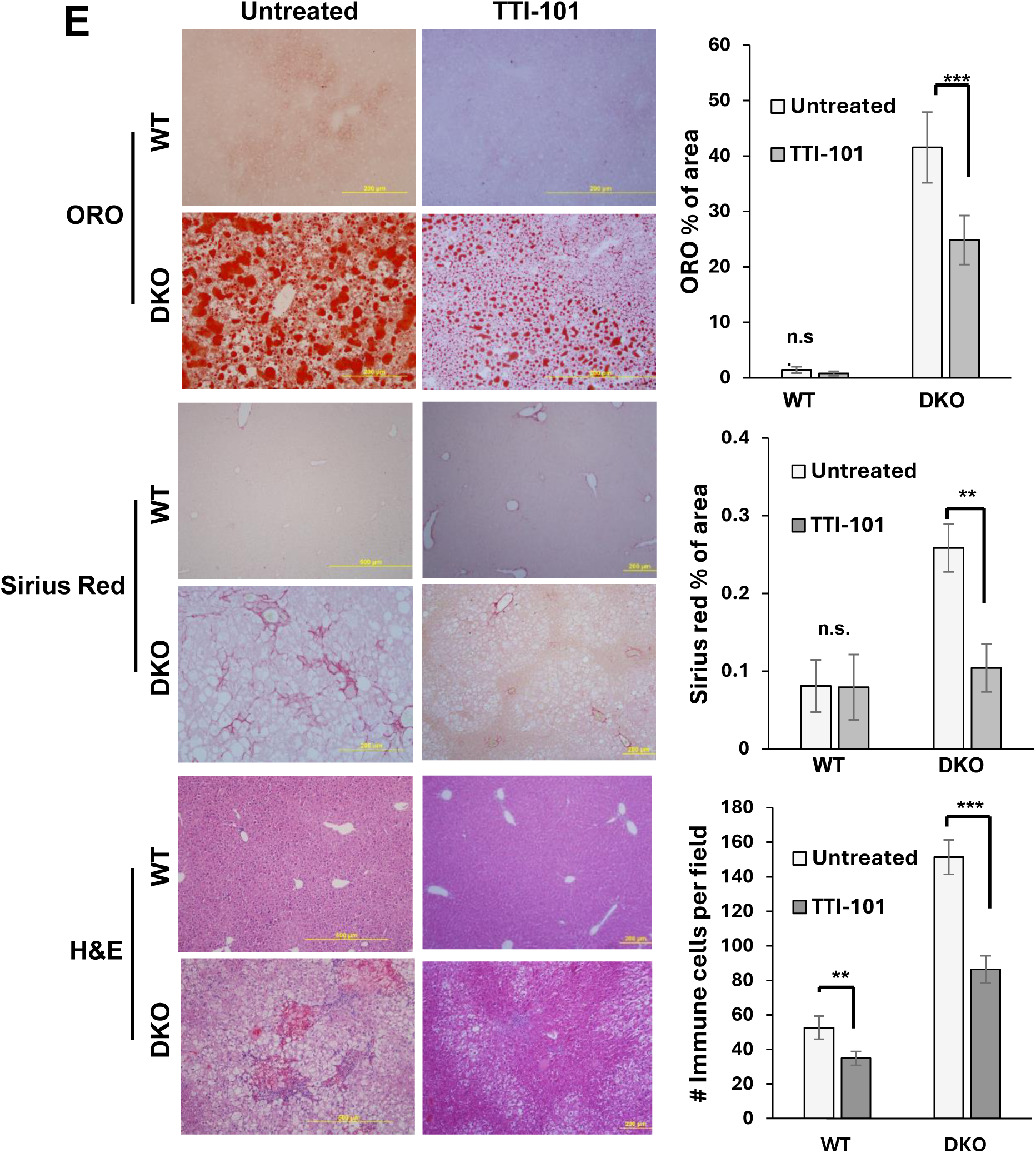
STAT3 is activated in hepatic stellate cells of DKO mice and its inhibition ameliorates MASH. **(A)** heatmap of mRNA expression of pro-inflammatory genes among the four genotypes. **(B)** WB analysis of pSTAT3-Y705, total STAT3 and Actin (control) in total liver lysates in each indicated genotype. **(C)** immunohistochemistry staining of α-SMA and pSTAT3-Y705 in consecutive DKO liver sections; the smaller dashed oval highlights the same landmark on each slide and the large oval highlights consecutive areas positive for both α-SMA and nuclear pSTAT3-Y705 (scale bar, 100 µm). **(D)** WB analysis of pSTAT3-Y705, total STAT3 and Vinculin (control) in untreated and TTI-101 DKO total liver lysates. **(E)** neutral lipid staining by Oil Red O (top), Sirius Red fibrosis (middle) and H&E inflammation analyses (bottom) in WT and DKO liver sections from untreated and TTI-101 treated mice, quantified to their right sides. (n.s.: not significant, **p<0.01, ***p<0.001 by 2-way ANOVA).

### 7. Activated STAT3 in HSCs accelerates the onset of MASLD-to-MASH progression in BRUCE and PTEN double-deficient mouse model of MASH

Due to the inadequate understanding of whether activated STAT3 in HSCs is responsible for driving its pro-fibrogenic function, we sought to address this critical question in our DKO mouse model of MASH using a STAT3-specific inhibitor TTI-101 (formerly C188-9). TTI-101 is a competitive inhibitor of STAT3 designed to target the phospho-tyrosine-peptide binding site within the SH2 domain of STAT3 and thereby blocks STAT3 activation^37,38^. TTI-101 was selected for this testing among other STAT3 small molecule inhibitors because it has no toxicity or interference with mitochondrial function in animal studies and clinical trials^39,40^. An earlier study has shown an ameliorating effect of TTI-101 on both MASH and HCC that are developed in PTEN-KO mice, yet there was no information provided to distinguish which liver cell type STAT3 is activated^41^. Therefore, we tested the effect of inhibiting STAT3 *in vivo*, by administration of 100mg/ kg TTI-101 to 2-month-old DKO and WT mice via daily intraperitoneal (i.p.) injection, or left untreated, for three weeks. Mice administered with TTI-101 for the duration did not show any noticeable health issues or side effects, which is the same as previously reported. Analysis of the liver showed that TTI-101 treatment significantly reduced hepatomegaly and the liver-to-body weight ratio in DKO mice, with no noticeable changes in WT livers (**Fig. S3A-B**). The inhibitor decreased both total and active STAT3 compared to untreated DKO mice in whole liver homogenates (**Fig. 6D**). TTI-101 effectively ameliorated lipid accumulation, liver fibrosis, and immune cell infiltration (**Fig. 6E**). These data show biochemical, and histopathological beneficial effects by the inhibition of pSTAT3-Y705. Collectively, TTI-101 shows both anti-MASLD and anti-MASH activities and that STAT3 activation in HCSs in BRUCE and PTEN DKO liver is a major driver for MASLD and MASH development.

### 8. *In vivo* specificity of TTI-101 on inhibition of HSC activation and the broad anti-MASLD and anti-MASH effects of TTI-101

To ascertain the *in vivo* specificity of TTI-101, we investigated whether it targets the activated STAT3 for inactivation in the HSC population in DKO mice. By IHC analysis of the respective liver tissue sections, we found that in TTI-101 treated DKO mice the number of activated HSCs were reduced compared to untreated DKO mice (**Fig. 7A**). In addition, the remaining activated HSCs (expressing a-SMA marker) were negative for nuclear expression of pSTAT3-Y705 (**Fig. 7B**). These results demonstrate a specific STAT3 inhibitory activity of TTI-101 in HSCs and the results also validate a driver function of activated STAT3 in HSCs in MASLD and MASH.

**Figure 7.**
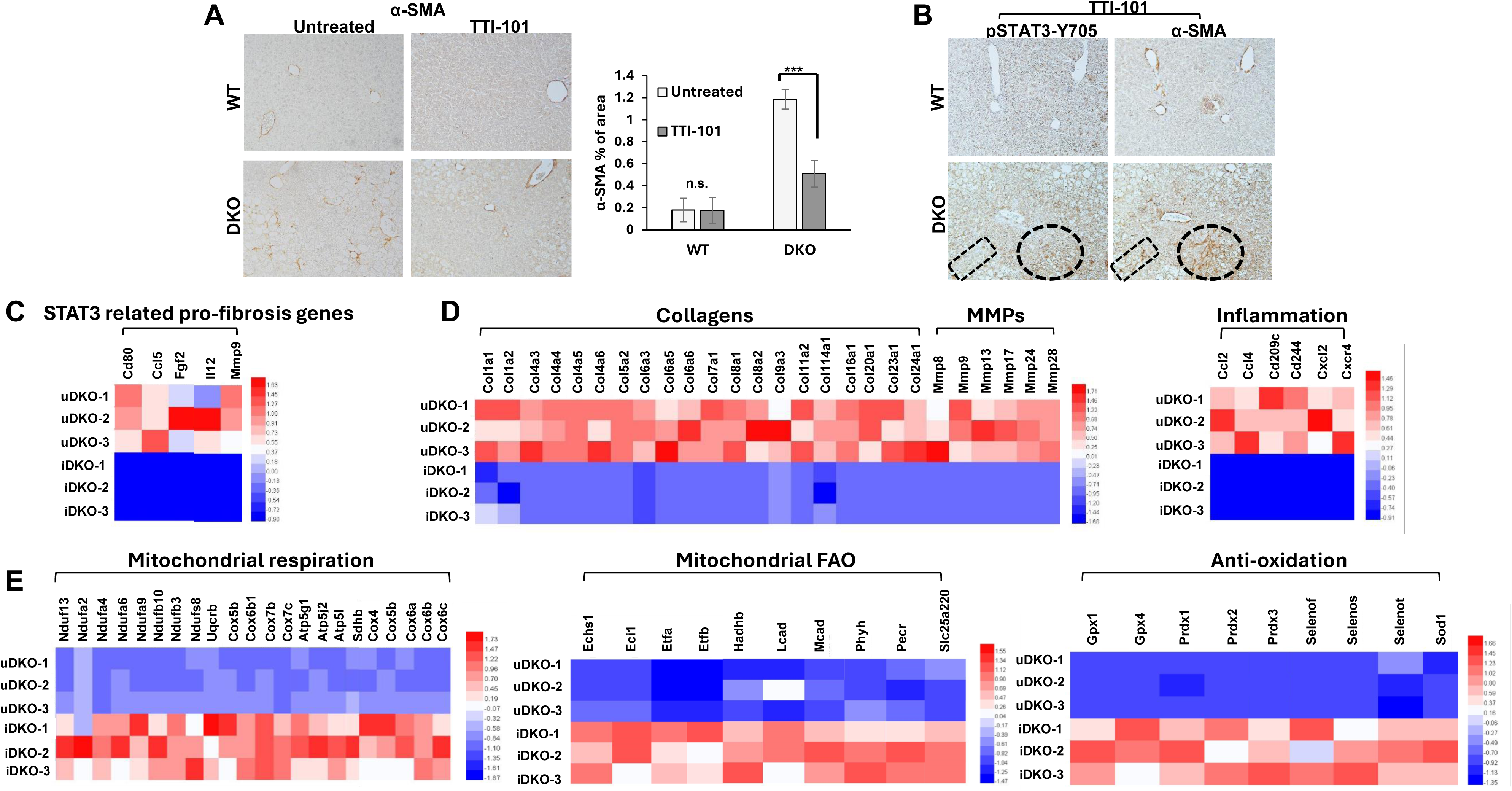
TTI-101 targets pSTAT3-Y705 in hepatic stellate cells to exert anti-MASLD/MASH effects. **(A)** α-SMA staining of activated hepatic stellate cells (HSCs) in WT and DKO untreated and TTI-101 treated mice. **(B)** immunohistochemistry analysis of pSTAT3-Y705 and α-SMA colocalization in consecutive slides from the indicated mice; the dashed rectangle and oval highlight α-SMA positive areas that are negative for nuclear pSTAT3-Y705 to show the same area on each slide. The RNA sequencing analysis generated heatmaps of relative mRNA expression of STAT3-related pro-fibrosis genes **(C),** collagens, MMPs and pro-inflammatory genes **(D)**, mitochondrial respiration and fatty acid oxidation (FAO) genes **(E)**, and genes involved in antioxidant systems **(F)**. (u=untreated, i=inhibitor treated, n.s.: not significant, ***p<0.001 by 2-way ANOVA).

To elucidate the mechanism by which TTI-101 resolves fibrosis and alleviates inflammation, we conducted whole liver transcriptomics with livers from DKO mice treated with or without TTI-101. As STAT3 transcriptionally activates a plethora of target genes, mostly identified in the field of STAT3 in tumorigenesis, we found TTI-101 effectively inhibited key target gene transcripts of pro-inflammatory cytokines (IL12, IFNγ, CCL5), M1 polarization (CD80), and matrix metalloproteinase 9 (MMP9), a main enzyme implicated in ECM degradation for remodeling the ECM, and regulating immune responses by facilitation of immune cell infiltration^42^ (**Fig. 7C**). The inhibitor also downregulated FGF2, an essential activator of STAT3 signaling^43^ (**Fig. 7C**), which could further turn down STAT3 signaling. To further elucidate the hepatic beneficial effects of TTI-101 attributed to its direct inhibition of STAT3 activity in HSCs, we found a significant downregulation of 20 key pro-fibrotic collagen gene transcripts, an equally important reduction of 6 pro-fibrotic MMP gene transcripts and reduction of inflammatory gene transcripts (**Fig. 7D**). Taken together, the findings of transcript downregulation demonstrate dual anti-fibrotic and anti-inflammatory activity of TTI-101 through switching off the fibrogenic and immunogenic signals to allow resolution of hepatic fibrosis and immune response in the DKO liver. In contrast to transcript downregulation, we found TTI-101 dependent, significant upregulation of transcripts involved in mitochondrial respiration, ATP production and mitochondrial FAO (**Fig. 7E**), demonstrating a pro-mitochondrial metabolic activity of TTI-101, negating the mitochondrial functional deficiency induced by BRUCE liver-KO. As oxidative stress plays critical roles in accelerating MASLD and MASH, administrated TTI-101 showed an activity of restoring hepatic antioxidant capacity in DKO livers by upregulation of multiple antioxidant enzyme transcripts critical for removal of oxidative stress. Both glutathione peroxidases 1 and 4 (GPX1 and GPX4) were upregulated and have the activity to reduce oxidized biomolecules at the expense of glutathione. The superoxide dismutase type 1 (SOD1) transcript was upregulated which is a copper/zinc superoxide dismutase that converts superoxide radicals into oxygen and hydrogen peroxide. In addition, transcripts of peroxidases of the peroxiredoxin family (PRDX1, 2, 4) were upregulated which reduce hydrogen peroxide and alkyl hydroperoxides to water and alcohol (**Fig. 7F**). These gene expression results agree with the beneficial effect of TTI-101 on reducing liver fibrosis and immune cell infiltration (**Fig. 6E**). Collectively, the findings of transcript upregulation demonstrate oxidative stress-lowing activity of TTI-101 to facilitate resolution of hepatic fibrosis and suppression of immune response in DKO liver.

### 9. Clinical relevance of dual downregulation of BRUCE and PTEN in association with STAT3 activation in healthy and MASH patient specimens

To gain insights into the clinical relevance of our findings from the above animal study, we performed IHC to examine the expression of BRUCE, PTEN and activated STAT3 in human MASH specimens (n = 4) with healthy human liver specimens as a control (n = 5). Images taken under a lower magnification (4X) showed a robust expression of both BRUCE and PTEN in the healthy liver and greatly reduced expression of both in MASH samples, supporting the involvement of their downregulation in MASH development (**Fig. 8A**). On the contrary, active STAT3 expression, indicated by pSTAT3-Y705, was lower in normal livers whereas higher in MASH livers, supporting the involvement of increased STAT3 activation in MASH development. A closer examination of their expression under a higher magnification (20X) showed decent expression of both BRUCE and PTEN in the hepatocytes of healthy livers whereas greatly reduced expression of both in MASH samples. In the areas of oval outlines, we observed downregulation of BRUCE and PTEN was concurrent with upregulation of activated STAT3 (pSTAT3-Y705) and the activated STAT3 was confined to the cell nucleus of non-hepatocyte areas, likely HSCs (**Fig. 8A-B**). Collectively, the data demonstrates a clinical correlation of dual downregulation of BRUCE and PTEN with upregulation of active pro-fibrotic STAT3 signaling in non-hepatocytes and mostly likely HSCs as suggested by our animal studies (**Fig. 6C**). As STAT3 activation in non-hepatocytes in patient livers has been shown to have a strong correlation with severity of liver fibrosis ^36^, our finding of HSCs bearing activated STAT3 suggests that these HCSs in the MASH patient subpopulation with BRUCE and PTEN dual downregulation is critical to the severity of their liver fibrosis.

**Figure 8.**
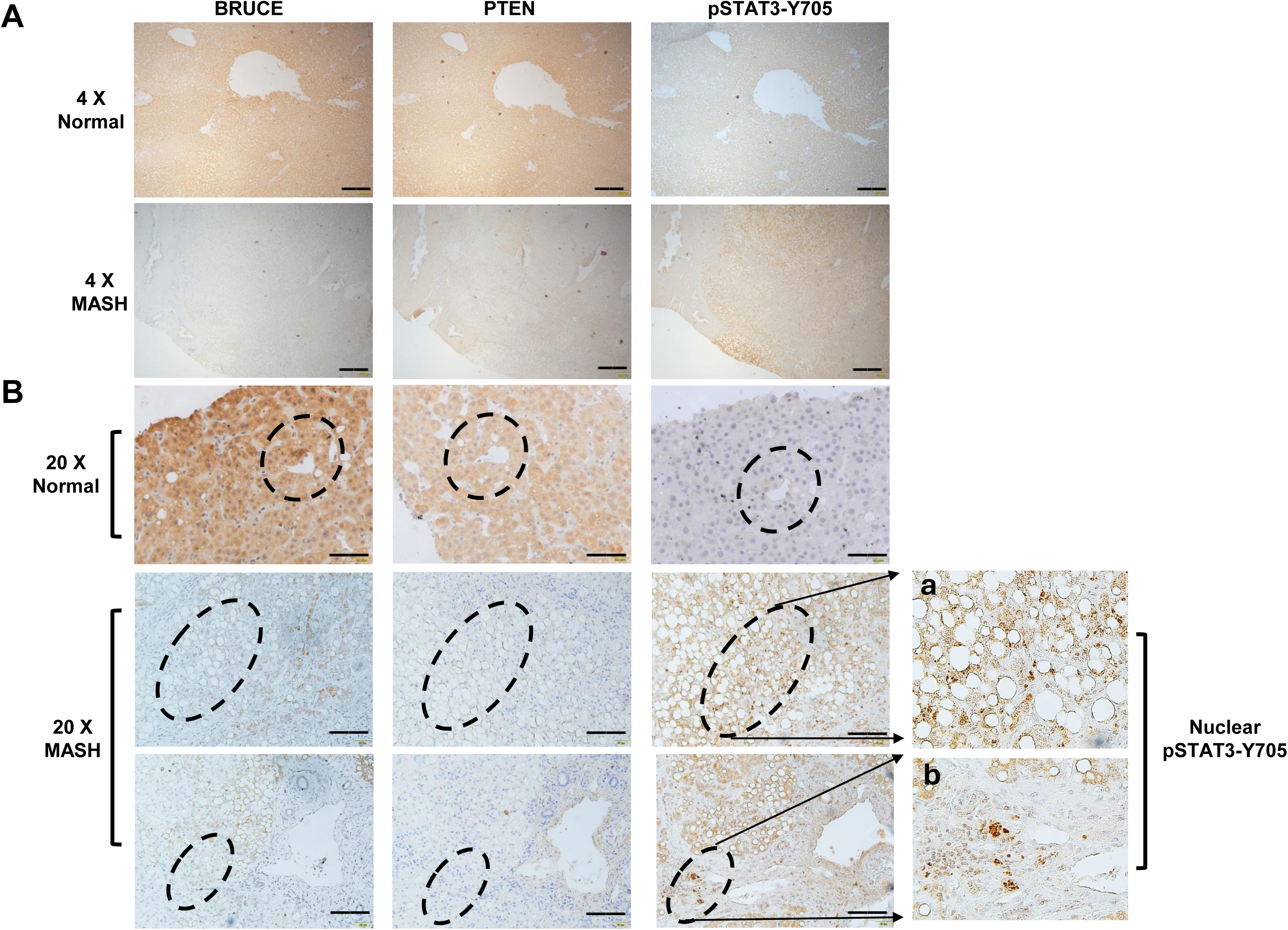
Clinical relevance of BRUCE and PTEN dual downregulation in MASH patient specimens. **(A)** 4x magnification of immunohistochemistry (IHC) analysis of BRUCE, PTEN and pSTAT3-Y705 expression in human normal (upper) and MASH specimens (lower) (scale bar, 400 µm). **(B)** IHC was performed on consecutive liver sections to show expression of BRUCE, PTEN and pSTAT3-Y705 in normal (upper) and MASH (lower) specimens in 20x magnification; the dashed ovals highlight landmarks to show the same area on each slide (scale bar, 100 µm).

## Discussion

This study of BRUCE and PTEN dual liver-KO mouse models provides new evidence that BRUCE and PTEN work in concert in the negative regulation of MASLD and MASH and BRUCE being a new co-suppressor of MASLD/MASH. We show that BRUCE-KO 2-month-old mice do not yet show any MASLD or MASH, whereas PTEN-KO at the same age show little initiation of lipid accumulation but do not show any MASH. However, 2-month-old DKO mice display significant mitochondrial metabolism defects which drive the accelerated development of MASLD, and the aberrant activation of pro-fibrotic STAT3 signaling in HSCs drives the accelerated MASLD-to-MASH transition. The potentiation of MASH by BRUCE liver-deficiency in PTEN-deficient background highlights a critical MASH suppressive role of BRUCE, at least through BRUCE’s role in maintaining mitochondrial lipid catabolism, respiration and bioenergetics. Further, this co-requirement for BRUCE and PTEN in the suppression of MASLD/MASH is a new finding. To effectively induce MASLD/MASH, BRUCE liver-deficiency that impairs mitochondrial function requires a strong lipogenic background induced by PTEN deficiency-mediated elevation of AKT-PI3K lipogenic pathway. Only a portion of patients with MASLD will develop MASH and the key determinants in this partial progression are not fully elucidated, yet this study indicates BRUCE deficiency may promote this transition.

The roles of BRUCE in apoptosis inhibition and DNA damage repair are well characterized, but its impact on hepatocyte mitochondrial metabolism had not been reported. The current study has provided *in vivo* and *in vitro* evidence of such a role in the context of MASLD/MASH. It is important to further investigate how BRUCE specifically regulates mitochondrial metabolism at molecular levels. Our earlier study reported a crucial effect of BRUCE knockdown on elevation of AMPK-mediated autophagy activation in human tumor cell lines, which links BRUCE to the regulation of cellular energy metabolism as AMPK is a crucial sensor of energy reduction^18^. As autophagy has a tight association with mitochondrial metabolism, the autophagic function of BRUCE in the development of MASLD and MASH is of pursuing in the future.

The PTEN regulation of lipogenesis through suppression of PI3K-AKT lipogenic signaling in mice is a well characterized mechanism for how PTEN liver-KO mice initiate MASLD at 3-months-old and full liver MASLD at 6-months-old, and eventually progress to liver cancer initiating around 12-months-old^5,9^. Now the current study adds BRUCE to PTEN context. PTEN also works in concert with multiple other factors in suppression of MASLD/MASH, one being the tyrosine phosphatase SHP2^44^. Further studies to examine the role of BRUCE in PTEN and SHP2 dual deficient background could provide new pathogenic mechanisms for these multifactorial liver diseases. The information could help patient stratification for individualized medication.

Although PTEN mutations is an infrequent event in hepatocytes^45,46^, decreased or absent PTEN protein expression is about 50% of primary hepatoma patients^47^. Decreased PTEN expression has a positive correlation with a higher tumor grade, advanced disease stage, and worse prognosis^47^. Liver fibrosis in PTEN liver-KO mice develops at a relatively later stage around 10 months of age. It is likely because MASLD/MASH is not a single factorial disease. BRUCE downregulation is found in a large percentage of MASLD/MASH and HCC specimens^15^, implying that BRUCE is a critical hepatic factor that dictates the disease progression in PTEN loss of function background. We have reported *BRUCE* somatic mutations are found in the TCGA HCC database^15^. These BRUCE somatic mutations could also play a role in the pre-HCC stage of MASLD/MASH. Further, some of these mutations coexist with PTEN mutations in the same individual although at a low frequency (data not shown) and individuals with the dual mutation could have a higher tendency to develop MASLD/MASH and progression to HCC. In contrast to somatic mutations, the much higher frequency of downregulation in BRUCE protein levels that we previously published in patient specimens^15^ and the reduction of both BRUCE and PTEN protein levels in all five MASH specimens presented in this study suggest loss/downregulation of both protein expressions could be another major liver disease promoting mechanism.

A study of 133 MASLD patients found that STAT3 activation in non-hepatocyte areas is strongly associated with fibrosis severity, inflammation, and progression to MASH^36^. Among many types of NPCs, HSCs are major collagen producing cells for fibrogenesis. Depletion of HCSs inhibits MASLD and MASH development in mice^48,49^. Our BRUCE and PTEN DKO mouse models also display activation of STAT3 in non-hepatocytes, HSCs, and the activated STAT3 in HSCs drives liver fibrosis, suggesting the BRUCE PTEN DKO mouse model mimics patient MASH features and the mechanism derived from the DKO mouse model could provide new solutions for MASH therapy. The in vivo ameliorative effect of STAT3 inhibitor TTI-101 in the DKO mouse model of MASLD/MASH indicates BRUCE liver-insufficiency should be a new crucial factor to be added to clinical consideration for clinical trials. In addition, hepatocytes stressed from excessive lipid accumulation release intracellular factors, cytokines, and growth factors, which bind to their receptors on HSCs thereby activating JAK-STAT3 pathway. These factors could be mitochondrial materials released from DKO hepatocytes because mitochondrial damage is a defining feature in DKO mice. Identification of these pro-inflammatory and pro-fibrogenic factors of mitochondrial materials in the future will provide new insights to the pathogenic mechanisms and potentially develop new therapeutic interventions suppressing inflammation and fibrosis in BRUCE and PTEN dual deficient MASH patients.

## Materials and Methods

### Animal Studies

All animal studies were performed in accordance with Institutional Animal Care and Use Committee approved protocols. The liver specific *Bruce* knockout (BKO) mice were generated as described previously^15^. To generate the liver specific *Pten* knockout (PKO) mice, *Pten*^flox/flox^ mice (The Jackson Laboratory; stock number 006440) were crossed with Alb-Cre mice (The Jackson Laboratory; stock number 018961). To generate the liver-specific *Bruce* and *Pten* double knockout (DKO) mice, BKO were crossed with PKO mice. Genotypes were confirmed by PCR. Littermates that are Cre negative were used as WT controls. Male mice were used in all experiments. For *in vivo* treatment, 2-month-old WT and DKO mice received daily intraperitoneal injections of 100mg/kg TTI-101 (MedChem Express, HY-112288) for three weeks.

### Liver Histology, Immunohistochemistry, TUNEL and DHE Staining

Mouse livers were removed and rinsed with phosphate buffered saline (PBS). For hematoxylin and eosin (H&E) staining, small pieces of liver were fixed in 10% neutral buffered formalin and embedded in paraffin and sectioned at 6 µm. Sections were stained with hematoxylin and eosin. For Oil Red O staining, liver tissues were embedded in OCT compounds and frozen on dry ice. The sections were cut at 10 µm and stained with freshly made Oil Red O solution (0.3% Oil Red O in 60% isopropanol). For Sirius Red staining, paraffin embedded liver sections (6 µm) were de-waxed and stained with Sirius Red solution (saturated picric acid containing 0.1% Sirius Red F3B).

For immunohistochemistry, paraffin embedded liver sections (6 µm) were de-waxed, rehydrated and antigen retrieved by using sodium citrate buffer (pH 6.5) or EDTA buffer (pH 8.0). After blocking endogenous peroxidase using 3 % H2O2, sections were then blocked with 5 % normal goat serum in PBS (containing 0.1% Triton X-100). Sections were incubated with primary antibodies overnight at 4 °C, after which Vectastain ABC kit (Vector Laboratories) was used, following manufacturer’s instructions. Sections were then incubated with 3,3′-Diaminobenzidine (DAB) and counterstained with hematoxylin. Terminal deoxynucleotidyl transferase dUTP nick end labeling (TUNEL) staining was performed with paraffin embedded sections using a TUNEL assay kit following the manufacturer’s instructions (Millipore, S7110). For DHE staining, frozen liver sections were stained with 2 µm DHE (Invitrogen, D11347) for 30 min at 37°C.

### Western Blot

Liver tissues were homogenized and lysed in RIPA buffer (20 mM Tris-HCI pH 8.0, 150 mM NaCl, 2 mM EDTA, 1% NP40, 1% sodium deoxycholate, 0.1% SDS) supplemented with Halt protease inhibitors (Thermo Scientific, 88668) and Pierce Universal Nuclease (Thermo Scientific, 88700). After clearing via centrifugation, the supernatant was collected for western blot analyses.

### RNA-Seq and Real-time qPCR

Liver RNA isolation was performed using the mirVana™ miRNA Isolation Kit (Thermo Scientific, AM1560) according to the manufacturer’s protocol. RNA-seq was performed at the University of Cincinnati Genomics, Epigenomics and Sequencing Core. Heatmaps were generated using the Heml 1.0 software. For Real-time PCR, RNA was reverse-transcribed using iScript cDNA Synthesis Kit (Bio-rad, 1708891). cDNA was then subjected to Real-time PCR analysis with SYBR Green Supermix (Bio-Rad, 1725121) in a CFX Connect Real-Time PCR Detection System (Bio-Rad, 1855201). Data were normalized to GAPDH control.

### Primary Hepatocyte Isolation

Mice were anesthetized and the inferior vena cava was cannulated with a 25-gauge cannula (Fisher Scientific 02-664-2), followed by incision of the portal vein. Livers were perfused and digested and hepatocytes isolated according to published method^50^ with cell viability assessed by trypan blue exclusion.

### Seahorse Analysis to Measure Mitochondrial Functions

Primary hepatocytes were isolated from 2-3-month-old male mice of each genotype. Hepatocytes were seeded in hepatocyte medium supplemented with 10% FBS, 2% penicillin-streptomycin, 1% sodium pyruvate, 1% L-glutamine, and 1% insulin-transferrin-selenium (Williams E Medium, Thermo Scientific, 12551032) at a density of 1 x 10^4^ cells per well in a Seahorse XF 96-well cell culture microplate (Agilent, 103794-100). After 24 hours, fresh hepatocyte medium was added, containing either BSA or palmitic acid (PA, Cayman Chemical, 10006627) conjugated to BSA. After 48 hours, the medium was changed to Seahorse assay medium supplemented with 10 mM glucose, 1 mM pyruvate, and 2 mM glutamine (Agilent 103575-100). The long chain fatty acid oxidation stress test (Agilent, 103672-100) was performed to measure oxygen consumption rates (OCR) using the Seahorse XFe96 Analyzer (Agilent, S7800B) according to the manufacturer’s protocol. OCR was measured after sequential injections of 4 µm etomoxir, 1.5 µm oligomycin, 1 µm FCCP, and 0.5 µm Rotenone/Antimycin A with 2.5 µm Hoechst. OCR values were normalized to cell number via fluorescence intensity of Hoechst staining (Fisher Scientific, 51-17).

### Data Analysis

Measurements were recorded in Excel (Microsoft). The results are expressed as the means ± SD or SEM of the determinations. ImageJ, CFX Connect (BioRad), Wave (Agilent Technologies), and CLARIOstar (BMG Labtech) softwares were used for data acquisition and quantification. Statistical significance was determined by the two-tailed Student’s *t*-test or 2-way ANOVA using Excel and GraphPad Prism with *p*<0.05. Bar graphs were generated in Excel.

## Supporting information

Che et al Supplemental Figures

Che et al Supplemental Figure Legends

Che et al Supplemental Methods

## Acknowledgments

This work was supported by the National Institutes of Health RO1 R01CA158323; R21 CA241025-01, the Center for Environmental Genetics, National Institutes of Environmental Health (NIH/NIEHS) grant P30-ES006096; Center for Clinical and Translational Science and Training (CCTST) Innovative Translational Pilot Award; University of Cincinnati College of Medicine Innovation Award. We thank our colleagues for constructive comments through the department seminar series and the MSTiC work in progress seminar.

