## Supplementary figures and images for "BRUCE liver-deficiency potentiates MASLD/MASH in PTEN liver-deficient background by impairment of mitochondrial metabolism in hepatocytes and activation of STAT3 signaling in hepatic stellate cells"

### Che et al Supplemental Figures

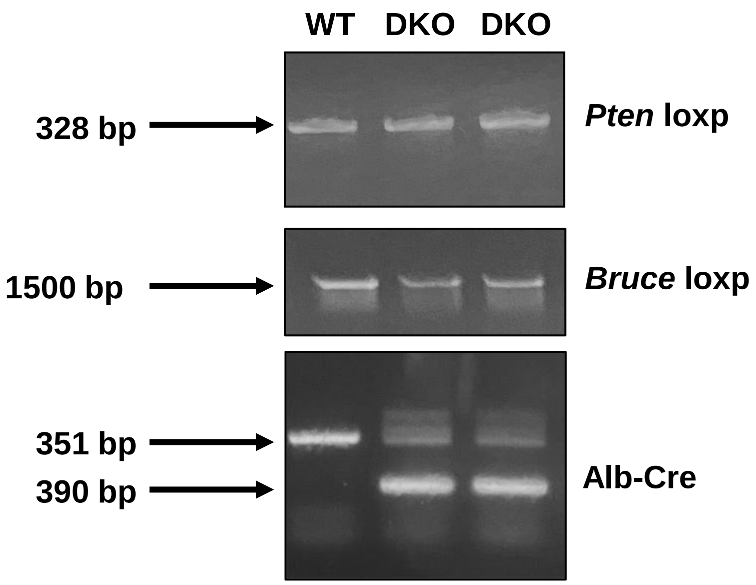

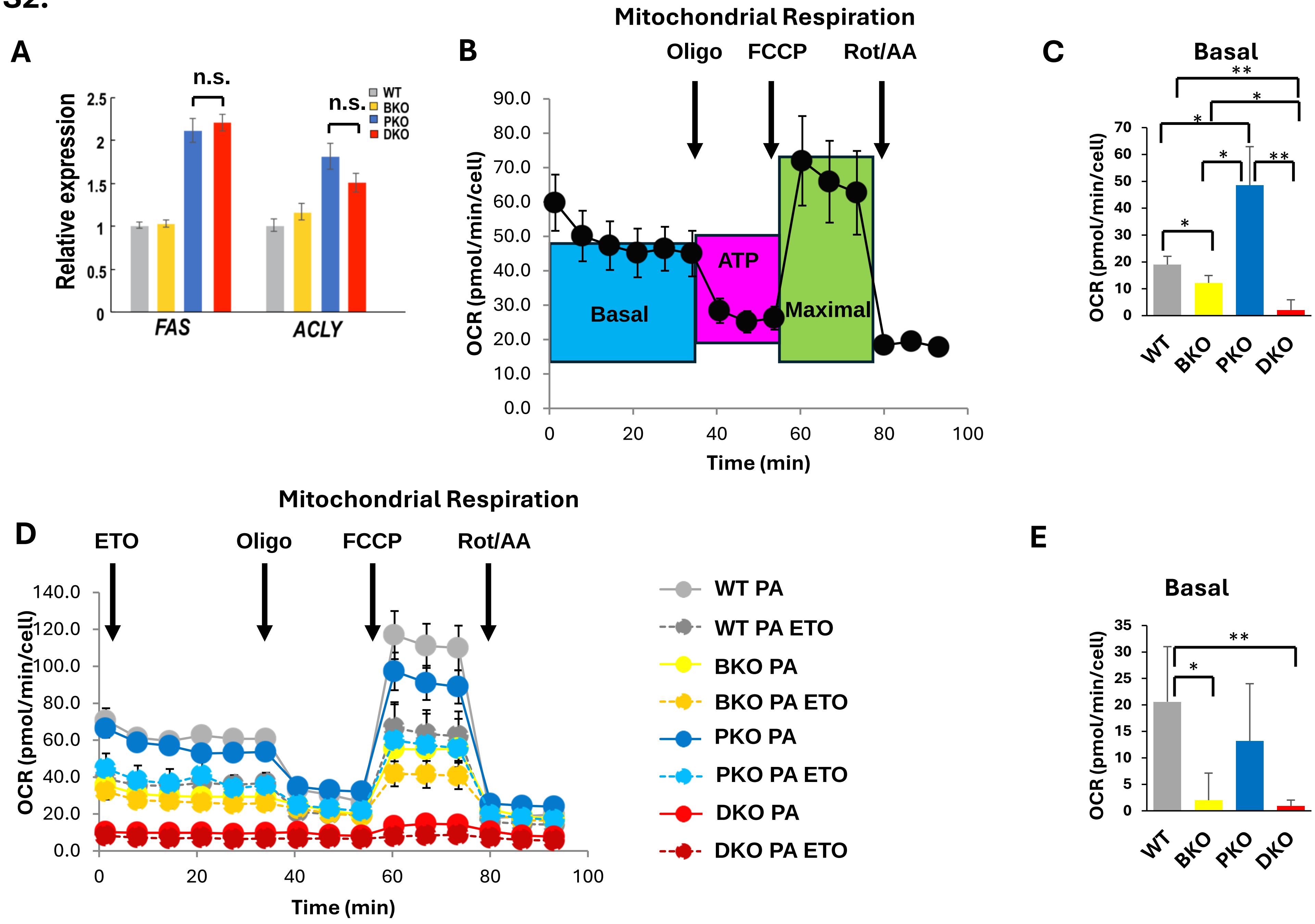

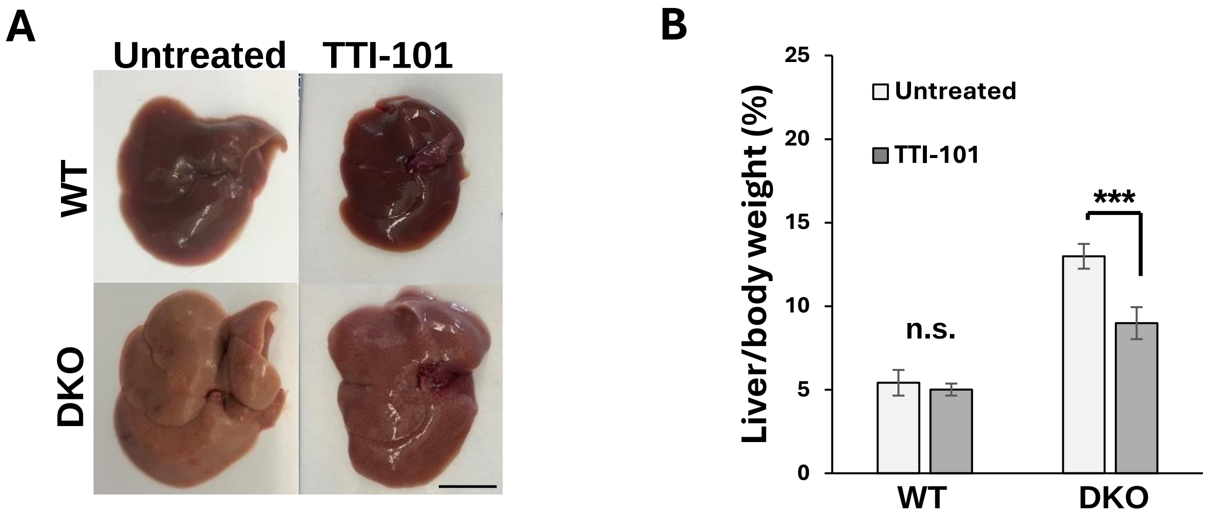
