## Supplementary material for "BRUCE liver-deficiency potentiates MASLD/MASH in PTEN liver-deficient background by impairment of mitochondrial metabolism in hepatocytes and activation of STAT3 signaling in hepatic stellate cells": Che et al Supplemental Figure Legends

**Figure S1. Genotyping BRUCE and PTEN double liver-KO mice.** PCR analysis of genomic DNA from mouse ear tissue generated from WT (*Bruce*$^{fl/fl}$*Pten*$^{fl/fl}$AlbCre-) and DKO (*Bruce*$^{fl/fl}$*Pten*$^{fl/fl}$AlbCre+).

**Figure S2. Genotyping BRUCE and PTEN double liver-KO mice. (A)** mRNA expression of lipogenesis genes *FASN* and *ACLY* determined by qRT-PCR. **(B)** mitochondrial respiration and ATP production was analyzed in primary hepatocytes by the Seahorse XF analyzer following sequential injections of oligomycin (Oligo), carbonyl cyanide-p-trifluoromethoxyphenylhydrazone (FCCP), and rotenone + Antimycin A (Rot/AA). **(C)** basal respiration shown by oxygen consumption rate (OCR) in each indicated genotype. **(D)** mitochondrial respiration kinetics in primary hepatocytes following 24-hour treatment of palmitic acid (PA) and injection of etomoxir (Eto) prior to analysis. **(E)** basal respiration supported by FAO across genotypes (PA ETO values subtracted from PA values). (n.s.: not significant, *$p<0.05$, **$p<0.01$, by student's t test).

**Figure S3. Beneficial effect of TTI-101 on phenotypic characteristics of MASLD/MASH. (A)** macroscopic images of livers from WT and DKO mice after 3 weeks of STAT3-Y705 inhibitor (TTI-101) treatment via daily injection (i.p.) or no treatment (scale bar, 1cm). **(B)** liver to body weight ratio in indicated mice. (n.s.: not significant, ***$p<0.001$, by 2-way ANOVA).
