## Supplementary material for "BRUCE liver-deficiency potentiates MASLD/MASH in PTEN liver-deficient background by impairment of mitochondrial metabolism in hepatocytes and activation of STAT3 signaling in hepatic stellate cells": Che et al Supplemental Methods

**Antibodies**

Cell Signaling: α-SMA (19245), Stat3 (9139), Phospho-Stat3-Y705 (9145), PTEN (9559), AKT (9272), Phospho-AKT-S473 (4060), GSK3β (9315), Phospho- GSK3β-S9 (9336), Actin (3700). Sigma: BRUCE (PLA0098, western blot), Tubulin (T9026). BD Biosciences: BRUCE (611193, immunohistochemistry)

**Genotyping Primers**:

| | sequence 5'→3' |
|---|---|
| *Bruce* | SC1: ATGTGCTGGGGTGGCTCATCAAC<br>4-LOX1: TCCTGACTGAGACACTGA ACTGG |
| *Pten* | oIMR9554 Forward: CAAGCACTCTGCGAACTGAG<br>oIMR9555 Reverse: AAGTTTTTGAAGGCA AGATGC |
| Albumin Cre | Wild Type Forward: TGCAAACATCACATGCACAC<br>Common: TTGGCCCCTTACCATAACTG<br>Mutant Forward: GAAGCAGAAGCTTAGGAAGATGG |

**qRT-PCR Primers**

| **Gene** | **Forward Sequence 5'→3'** | **Reverse Sequence 5'→3'** |
|---|---|---|
| FASN | GCTGCGGAAACTTCAGGAAAT | AGAGACGTGTCACTCCTGGACTT |
| ACLY | GGTGTCAACGAACTGGCGAA | GTTTGCAATGCTGCCTCCAA |
| GAPDH | CAAAATGGTGAAGGTCGGTGTG | TGATGTTAGTGGGGTCTGGCTC |

**Hepatocyte Isolation Buffers**

Perfusion medium: Double-distilled water, 5.4 mM KCl, 0.3 mM Na2HPO4, 0.4 mM KH2PO4, 4.2 mM NaHCO3, 137 mM NaCl, 10 mM glucose, 10 mM HEPES, and 0.3 mM EDTA. pH adjusted to 7.4 and sterilized by filtration through a 0.22-micron filter.

Digestion medium: Double-distilled water, 5.4 mM KCl, 0.3 mM Na2HPO4, 0.4 mM KH2PO4, 4.2 mM NaHCO3, 1.3 mM CaCl2, 0.5 mM MgCl2, 0.6 mM MgSO4, 137 mM NaCl, 10 mM glucose, and 10 mM HEPES. pH adjusted to 7.4 and sterilized by filtration through a 0.22-micron filter. 0.05% collagenase IV prepared (Sigma C5138) and sterilized.
